# Catalytic-independent functions of INTAC in conferring sensitivity to BET inhibition

**DOI:** 10.1101/2024.02.07.579305

**Authors:** Pengyu Fan, Xue-Ying Shang, Aixia Song, Shuo Chen, Run-Yuan Mao, Jiwei Chen, Zhenning Wang, Hai Zheng, Bolin Tao, Wei Xu, Wei Jiang, Huijuan Yang, Congling Xu, Peng Zhang, Hai Jiang, Fei Xavier Chen

## Abstract

Chromatin and transcription regulators are critical to defining cell identity through shaping epigenetic and transcriptional landscapes, with their misregulation being closely linked to oncogenesis. Pharmacologically targeting these regulators, particularly the transcription activating BET proteins, has emerged as a promising approach in cancer therapy, yet intrinsic or acquired resistance frequently occurs with poorly understood mechanisms. Using genome-wide CRISPR screens, we find that BET inhibitor efficacy in mediating transcriptional silencing and growth inhibition depends on the auxiliary module of the INTAC complex, a global regulator of polymerase pause-release dynamics. This process bypasses a requirement for INTAC’s catalytic activities and instead leverages direct engagement of the auxiliary module with the RACK7/ZMYND8–KDM5C complex to remove histone H3K4 methylation. Targeted degradation of the COMPASS subunit WDR5 to attenuate H3K4 methylation restores sensitivity to BET inhibitors, highlighting how simultaneously targeting coordinated chromatin and transcription regulators can circumvent drug-resistant tumors.

## INTRODUCTION

Chromatin and transcription regulators are foundational elements for the establishment of cell identity through the orchestration of epigenetic and transcriptional landscapes, while alterations of these landscapes can trigger oncogenic gene expression. However, these changes can render cancer cells uniquely dependent on certain regulatory proteins for survival, presenting a therapeutic opportunity of targeting these addictions.^1–5^ The bromodomain and extraterminal domain (BET) family of proteins, comprised of BRD2, BRD3, BRD4, and the testis-specific BRDT, has come under scrutiny for its role in regulating the expression of genes pivotal for cancer cell growth and survival. BET family inhibitors, which target a conserved bromodomain, were designed to prevent these proteins from binding to chromatin, have emerged as promising anti-proliferative agents in cancer treatment. However, the clinical efficacy of these inhibitors is often limited by the occurrence of resistance to therapy, both intrinsic and acquired during treatment. Unraveling these resistance mechanisms is challenging but essential for optimizing the efficacy of BET inhibitors and advancing cancer treatment strategies.^6–23^

The epigenetic landscape is intricately shaped by the activity of enzymes that dynamically add or remove chemical modifications to histones or DNA. These enzymes, known as “writers” and “erasers” respectively, work in concert or opposition to modify chromatin structure and function. In addition to regulating chromatin compaction and nucleosome dynamics, these modifications are pivotal in regulating the activity of various chromatin and transcription regulators possessing “reader” domains that recognize their corresponding epigenetic marks.^24–27^ Histone acetylation, particularly at H3 lysine 27 (H3K27ac), is a hallmark of transcriptionally active chromatin found at promoters and enhancers.^5,28,29^ BET proteins can bind to the acetylated chromatin via their bromodomains where they can serve as a platform for recruiting other factors and help maintain acetylation levels established by acetyltransferases such as CBP and P300. ^15,30–35^ Histone acetylation is a highly dynamic and reversible histone mark that is removed by histone deacetylases (HDACs). Dominant among these are HDAC1 and HDAC2, which are integral components of several transcription corepressor complexes, including the nucleosome-remodeling HDAC (NuRD) complex, corepressor for REST (CoREST), and the SIN3 complex.^36–38^

Promoters and enhancers, the essential DNA elements for transcriptional control, are typically characterized by specific histone H3 lysine 4 (H3K4) methylation patterns—with relative enrichment of tri-methylation (H3K4me3) at promoters and mono-methylation (H3K4me1) for enhancers.^5,39,40^ The deposition of H3K4 methylation is mediated by the COMPASS family, which includes six members in mammals: SET1A, SET1B, MLL1, MLL2, MLL3, and MLL4. These enzymes function within complexes that share four core components—WDR5, RBBP5, ASH2L, and DPY30—collectively known as “WRAD”.^41–46^ Conversely, the removal of H3K4 methylation is catalyzed by two families of demethylases, the KDM1 family (LSD1/KDM1A and LSD2/KDM1B), which specifically removes mono- and di-methylation, and the KDM5 family (KDM5A to KDM5D), which preferentially targets di- and tri-methylated H3K4.^47–50^

Recent research has unveiled the dynamic interplay between COMPASS-mediated methylation and its removal by KDM5, with both processes exhibiting surprising flexibility.^51,52^ As with HDACs, these histone demethylases are components of various multi-subunit complexes. For example, LSD1 is part of the CoREST and NuRD complexes, which also have histone deacetylation activity, while LSD2 specifically form a complex with NPAC/GLYR1 targeting active gene body due to its affinity for H3K36me3.^53–55^ The KDM5 family can be incorporated into a RACK7/ZMYND8-containing complex and the SIN3 deacetylase complex. The RACK7-containing demethylase complex is equipped with multiple zinc-finger motifs and domains for binding histone modifications, which might influence its genomic distribution.^56–59^ However, the precise mechanisms determining the recruitment of the RACK7-containing complex to specific chromatin regions remain unclear. Furthermore, the intricate relationship between histone acetylation and methylation statuses, mediated by these multifaceted complexes, plays a crucial role in either promoting or repressing transcription under various biological contexts. This layered regulation highlights the need for further investigation into how these epigenetic marks and their modifiers collaborate to influence gene expression and cellular phenotypes.

As an essential regulator for virtually all classes of RNA polymerase II (Pol II)-transcribed genes, the Integrator–PP2A (INTAC) complex harbors both an RNA cleavage activity from Integrator and a phosphatase activity from the core PP2A (PP2A-A and PP2A-C).^60–63^ These two catalytic modules are structurally connected by the scaffolding backbone and shoulder modules, yet have different roles in transcription regulation at gene promoters or enhancers.^64–67^ Specifically, the RNA endonuclease module primarily induces promoter-proximal termination at promoters and attenuates non-productive transcription at enhancers, while the phosphatase module mainly prevents the release of paused Pol II to limit transcriptional activation.^66,68–70^ Moreover, Integrator and the larger INTAC assemblage contain a four-subunit (INTS10, INTS13, INTS14, and INTS15) auxiliary module, also known as the arm module, which is the least understood of the INTAC modules. Although the auxiliary module has an accessory role in INTAC endonuclease-mediated snRNA processing, with its loss resulting in a slight snRNA termination defect, subunits of this module may possess functions independent of Integrator or INTAC.^71–74^ For example, INTS13 was shown to be able to form an independent sub-module with transcription factors that targets poised enhancers to induce monocytic differentiation.^75^

Here, to explore the mechanisms through which BET inhibitors exert their effects in halting cancer cell proliferation and to understand how the intrinsic and acquired resistance can occur, we have leveraged genome-wide CRISPR screens to find that loss of the INTAC auxiliary module results in a profound resistance to BET inhibition. In contrast, the disruption of other INTAC modules, including the two catalytic modules, does not elicit alterations in sensitivity to BET inhibition, implying an independent role of the INTAC auxiliary module in mediating the response to BET inhibitors. We find that depletion of the auxiliary module impedes BET inhibitor-mediated transcriptional repression and the associated diminishment of epigenetic marks, including H3K4 methylation. Mechanistically, the INTAC auxiliary module directly interacts with the RACK7–KDM5C complex to facilitate its recruitment to chromatin and subsequent H3K4 demethylation. Moreover, we show that targeted degradation of the COMPASS subunit WDR5 effectively abates the aberrant accumulation of H3K4 methylation resulting from disruption of the INTAC auxiliary module and thereby resensitizing these tumor cells to BET inhibition.

## RESULTS

### CRISPR screens identify the INTAC auxiliary/arm module as a primary regulator of BET inhibition sensitivity

To elucidate the mechanisms conferring resistance to BET protein inhibition, we performed genome-wide CRISPR screens on lymphoma cells derived from Eμ-Myc transgenic mice, which were engineered to stably express Cas9.^76,77^ We introduced these cells to a lentiviral library of containing 188,509 sgRNAs targeting 18,850 protein-coding genes, as well as 199 non-targeting sgRNAs as negative controls. Following puromycin treatment to eliminate non-infected cells, two rounds of selection using BET protein inhibitor JQ1 were conducted to enrich cells with acquired resistance to BET inhibition (Fig. 1A). High-throughput sequencing of sgRNAs revealed that 32 genes were more than fourfold enriched in the JQ1-treated cells compared to the DMSO-treated controls (Fig. S1A). This group includes several genes implicated in the autophagy pathway, consistent with prior research suggesting a link between BRD4 depletion to the induction of autophagy-induced cell death (Fig. 1B and S1A, green).^78^

**Figure 1.**
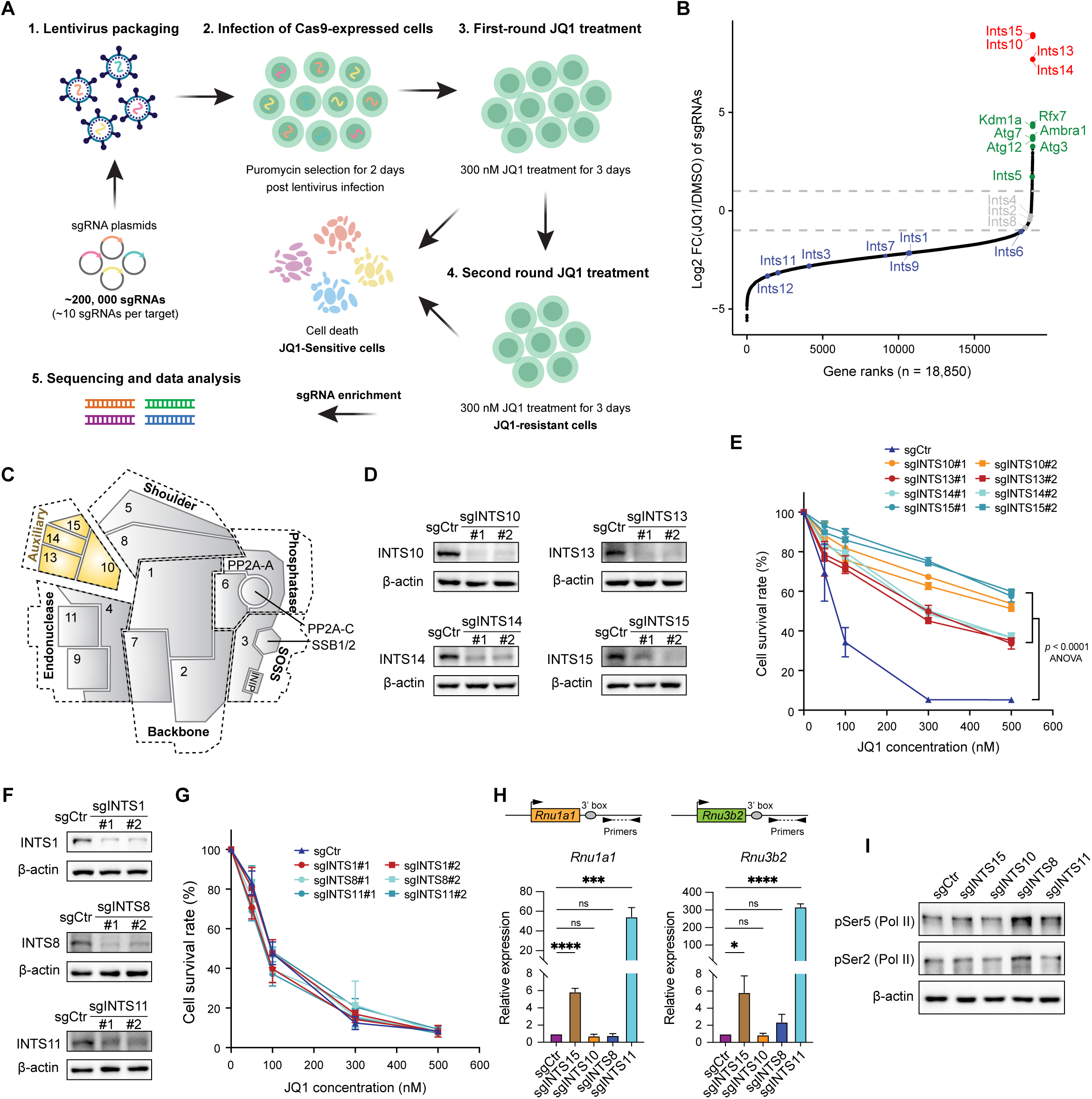
The INTAC auxiliary/arm module regulates BET inhibition sensitivity independently of INTAC’s catalytic activities. (A) Flow chart of the genome-wide CRISPR screening conducted to enrich cells with acquired resistance to BET inhibition in Eμ-Myc cells with the stable expression of Cas9. (B) Rank plot showing the log2 fold change (JQ1 versus DMSO) of sgRNA reads in survived cells post JQ1 treatment. Top ten enriched sgRNA targets and genes encoding INTAC complex labeled in color. (C) Schematic of the INTAC complex, components of the auxiliary module are highlighted. (D) Western blotting for INTS10, INTS13, INTS14 and INTS15 in CRISPR knockout Eμ-Myc cells. β-actin is a loading control. (E) Cell survival assays in sgCtr, sgINTS10, sgINTS13, sgINTS14 and sgINTS15 Eμ-Myc cells treated with DMSO or JQ1. (F) Western blotting for INTS1, INTS8 and INTS11 in CRISPR knockout Eμ-Myc cells. (G) Cell survival assays in sgCtr, sgINTS1, sgINTS8 and sgINTS11 Eμ-Myc cells treated with DMSO or JQ1. (H) Measurement for transcription termination of the representative snRNAs *Rnu1a1* and *Rnu3b2* in sgCtr, sgINTS15, sgINTS10, sgINTS8 and sgINTS11 Eμ-Myc cells. (I) Western blotting for RNA Pol II phosphorylation levels at carboxyl-terminal domain (CTD) Serine 5 (pSer5) and Serine 2 (pSer2) in sgCtr, sgINTS15, sgINTS10, sgINTS8 and sgINTS11 Eμ-Myc cells. β-actin is a loading control. See also Figure S1.

Of particular interest, four genes—*Ints15*, *Ints10*, *Ints13*, *Ints14*—were found to be over 100-fold enriched in our CRISPR screens (Fig. 1B, red). These genes encode all four subunits of the auxiliary module, also known as the arm module, of the Integrator or INTAC complex, which plays a pivotal role in transcriptional regulation of both protein-coding and non-coding genes (Fig. 1C).^79,80^ INTAC comprises at least six distinct modules, including two catalytic modules—an endonuclease module that cleaves various RNA species and a phosphatase module that dephosphorylates RNA Pol II and other substrates.^60–62^ The backbone and shoulder modules are essential for the assembly and, consequently, the catalytic activities of the complex.^60^ The SOSS module is known for recognizing single-stranded DNA and facilitating the recruitment of INTAC to R-loop-enriched regions.^81^ Nonetheless, the role of the auxiliary module in transcription remains less defined and it may harbor INTAC-independent functions (Fig. 1C).^75^

To validate our screening results, we individually depleted each of the four subunits of the INTAC auxiliary module subunits using two distinct sgRNAs in the pooled mouse lymphoma cells (Fig. 1D). Eliminating any one of these subunits led to an increase in resistance to the BET inhibitor JQ1 in a dose-responsive manner, with the removal of INTS15 and INTS10 showing the most substantial resistance (Fig. 1E). Furthermore, cells without the auxiliary module exhibited diminished sensitivity to BET inhibition by time-course JQ1 treatments (Fig. S1B). To further explore the relationship between the INTAC auxiliary module with the BET inhibition sensitivity, we compared mRNA levels of this module in human cancer cell lines from Cancer Cell Line Encyclopedia (CCLE)^82^ and their sensitivity to JQ1 inhibition, noting relatively lower expression of auxiliary subunits in JQ1 resistant cells (Fig. S1C–S1F). Furthermore, the targeted disruption of each auxiliary subunit in two randomly selected cancer cells of different types also induced the resistance to JQ1, suggesting a widespread role of the INTAC auxiliary module in modulating sensitivity to BET inhibition (Fig. S1G–S1H).

The exclusive enrichment of sgRNAs targeting the auxiliary module suggests it may uniquely influence JQ1 sensitivity, independent of other INTAC modules, especially the catalytic ones. To test this hypothesis, we performed targeted knockouts of INTS1 (a core component of the backbone module), INTS8 (a shoulder module subunit essential for phosphatase module recruitment), and INTS11 (the enzymatic subunit of the RNA endonuclease module) (Fig. 1C and 1F). Depleting these subunits did not markedly affect cellular sensitivity to JQ1 (Fig. 1G), reinforcing our CRISPR screening results.

In contrast to the severe readthrough in snRNA transcription observed in INTS11-depleted cells, the absence of INTS15 and INTS10 led to mild defects in snRNA processing (Fig. 1H), consistent with previous studies.^71–74^ Moreover, cells with INTS15 or INTS10 depletion did not show noticeable changes in Pol II phosphorylation observed in INTS8-depleted cells, which disrupts INTAC phosphatase modules and thus induces Pol II hyperphosphorylation (Fig. 1I).^62,66,83^ These findings suggest that the role of the auxiliary module extends beyond the primary catalytic functions of INTAC, possibly contributing independently to transcriptional regulation.

### The INTAC auxiliary/arm module facilitates BET inhibition-induced transcriptional repression

The BET protein family comprises four members, including BRD2, BRD3, BRD4, and the testis-specific BRDT, all of which are targeted by BET inhibitors like JQ1 (Fig. 2A). To identify the specific BET protein responsible for JQ1’s growth-inhibitory effects on cancer cells, we individually knocked out BRD2, BRD3, and BRD4 with two distinct sgRNAs in pooled cell populations (Fig. S2A). Notably, only the removal of BRD4 significantly hindered lymphoma cell proliferation, in contrast to the knockouts of BRD2 and BRD3 (Fig. 2B). We thus depleted INTS15 and INTS10 in BRD4-deficient cells, finding that either one destabilized each other (Fig. 2C) and markedly reversed the proliferation slowdown caused by BRD4 loss (Fig. 2D).

**Figure 2.**
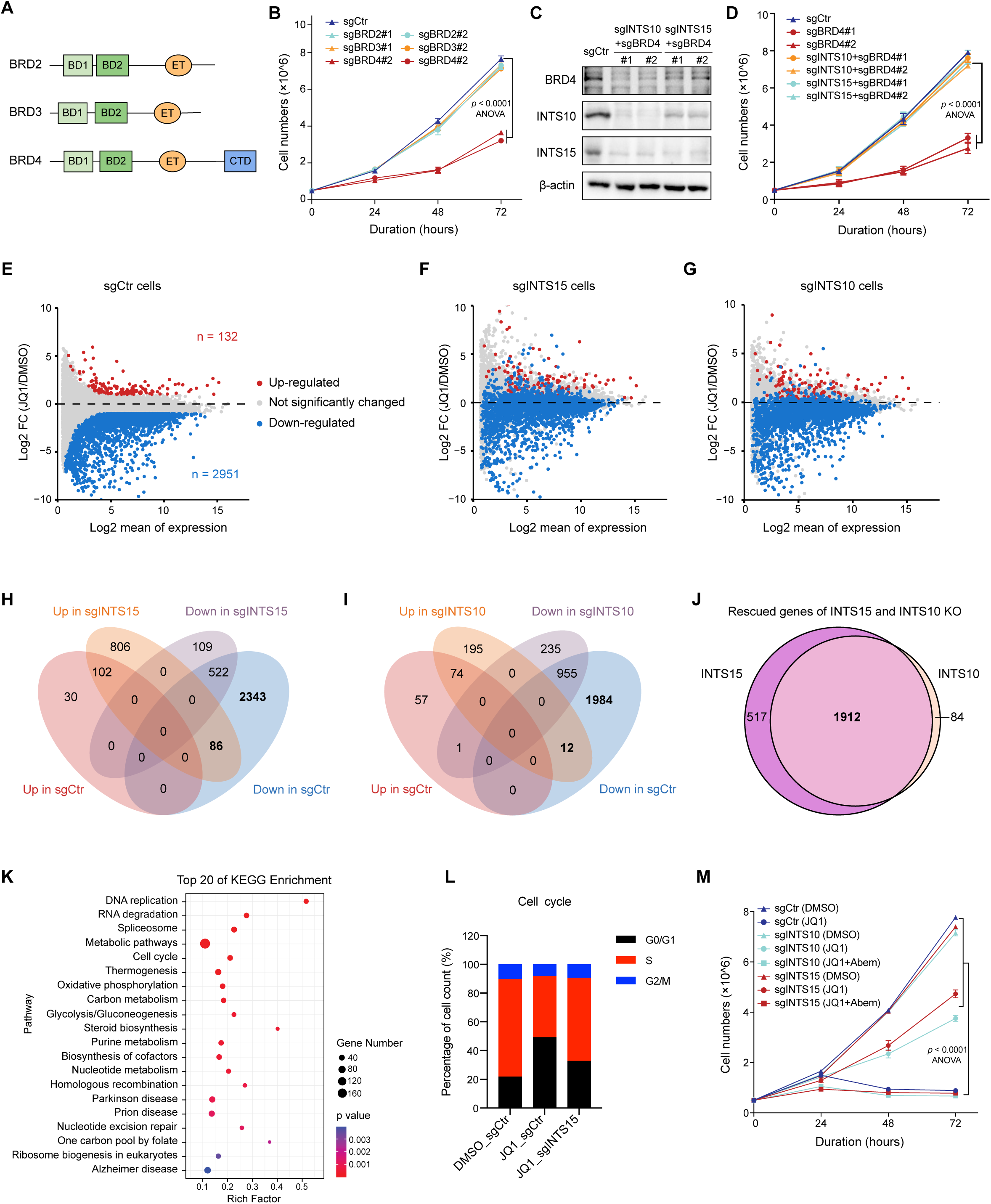
The expression of BET inhibition facilitates BET inhibition-induced transcriptional repression and cell cycle arrest. (A) Schematic showing the structures of BRD2, BRD3 and BRD4 in BET protein family. (B) The comparison on cell proliferation among sgCtr, sgBRD2, sgBRD3 and sgBRD4 Eμ-Myc cells. (C) Western blotting of BRD4, INTS10 and INTS15 in sgCtr, sgBRD4 as well as sgBRD4 co-transfected with sgRNA plasmid for INTS10 (sgBRD4; sgINTS10) or INTS15 (sgBRD4; sgINTS15) Eμ-Myc cells. (D) The comparison on cell proliferation among sgCtr, sgBRD4, sgBRD4; sgINTS10 and sgBRD4; sgINTS15 Eμ-Myc cells. (E) MA plot illustrating the up- and down-regulated genes in sgCtr Eμ-Myc cells post JQ1 treatment. (F and G) MA plot illustrating the up- and down-regulated genes in sgINTS15 (F) and sgINTS10 cells (G) post JQ1 treatment, in comparison to deferentially expressed genes (DEGs) in sgCtrl cells against JQ1 which labeled in dots. (H and I) Venn diagrams showing overlaps for up- and down-regulated genes in sgCtr and sgINTS15 (H) / INTS10 (I) Eμ-Myc cells post JQ1 treatment. The number of genes down-regulated in JQ1-treated sgCtr cells, but not repressed in JQ1-treated sgINTS15 / sgINTS10 cells labeled in bold. (J) Venn diagrams showing overlaps for genes down-regulated in JQ1-treated sgCtr cells, but not repressed both in JQ1-treated sgINTS15 and sgINTS10 cells. (K) KEGG pathway enrichment analysis for genes down-regulated by JQ1 but restored in JQ1-treated sgINTS15 and sgINTS10 cells. (L) Stacked plot showing the proportions of sgCtrl and sgINTS15 cells treated with DMSO or JQ1 across different phases of cell cycle. (M) Cell proliferation comparison among sgCtr, sgINTS10 and sgINTS15 cells treated with DMSO, JQ1 and combined JQ1 with CDK4/6 inhibitor Abemacialib. See also Figure S2.

Considering the role of as a transcriptional activator, we speculated that the INTAC auxiliary module regulates BET inhibition sensitivity by modulating gene expression. To examine this hypothesis, we conducted spike-in normalized RNA sequencing (RNA-seq) to evaluate JQ1-induced transcriptional changes. Principal component analysis (PCA) revealed that, while the depletion of INTS15 and INTS10 in DMSO-treated cells led to minor transcriptomic shifts, their removal in JQ1-treated cells caused significantly greater alterations (Fig. S2B). We then quantified the differentially expressed genes (DEGs) between DMSO and JQ1 treatment across the cell lines. In control cells, JQ1 treatment predominantly led to gene expression reduction, with 2951 genes significantly downregulated and 132 upregulated (Fig. 2E). Conversely, in cells lacking INTS15 and INTS10, JQ1 treatment elicited more moderate transcriptional changes, with 631 and 1191 genes significantly downregulated, respectively (Fig. S2C–S2D). Depicting these DEGs obtained from control cells onto MA plots for INTS15- and INTS10-null cells highlighted a clear mitigation of JQ1’s transcription repression upon depletion of these INTAC auxiliary subunits (Fig. 2F–2G). Analysis of these JQ1-induced DEGs suggested that 2429 (82.3%) and 1996 (67.6%) downregulated genes were rescued by the loss of INTS15 and INTS10, respectively (Fig. 2H–2I). Moreover, these rescued genes by the loss of INTS15 and INTS10 were largely overlapped, indicating their regulation is dependent on the INTAC auxiliary module (Fig. 2J).

### The INTAC auxiliary/arm module is required for BET inhibition-induced cell cycle arrest

Subsequent KEGG pathway analysis on the genes downregulated by JQ1 yet rescued by the loss of both INTS15 and INTS10 revealed significant enrichment in pathways related to DNA replication, RNA degradation and splicing, cell cycle regulation, and various metabolic processes (Fig. 2K), with several cell cycle related genes confirmed by the reverse transcription-quantitative PCR (RT-qPCR) (Fig. S2E). This led us to further explore the influence of JQ1 and the INTAC auxiliary on cell cycle progression. JQ1 treatment induced G1 phase arrest, characterized by an increased percentage of cells in G1 and a corresponding decrease in the S phase (Fig. 2L and S2F). Intriguingly, the elimination of INTS15 effectively mitigated the JQ1-induced cell cycle arrest (Fig. 2L and S2F). We thus posited that disrupting the G1 to S phase transition could counteract the resistance to JQ1 conferred by the absence of the INTAC auxiliary module. Confirming this hypothesis, inhibiting CDK4/6 with Abemacialib in cells lacking INTS15 and INTS10 increased the sensitivity to JQ1 in a dose-responsive manner (Fig. 2M and S2G). This suggests that the loss of the INTAC auxiliary module antagonizes BET inhibition-induced cell cycle arrest.

### The broad impact of the INTAC auxiliary/arm module in gene expression upon BET inhibition

To explore the mechanism by which the INTAC auxiliary confers BET inhibition-mediated transcriptional repression, we examined the expression changes by JQ1 treatment in INT15- and INTS10-depleted cells, ranking in the same order with the differential expression pattern observed in the control cells. Although the general trend of transcriptional alterations was consistent across the three cells, genes that were not significantly affected in control cells exhibited distinct responses to JQ1 in cells with INT15 and INTS10 depletion, implying the broad impact of the INTAC auxiliary on transcription (Fig. S3A– S3C). Notably, the absence of INTS15 and INTS10 resulted in a general increase in gene expression levels, regardless of their response to JQ1 (Fig. 3A). Indeed, by grouping genes into quartiles based on their expression changes due to JQ1, we observed that the deletion of INTS15 and INTS10 not only mitigated the transcriptional suppression induced by JQ1 but also enhanced the expression of genes irresponsive to JQ1 (Fig. 3B and S3D). These findings underscore the role of the INTAC auxiliary in attenuating gene expression in the context of BET inhibition.

**Figure 3.**
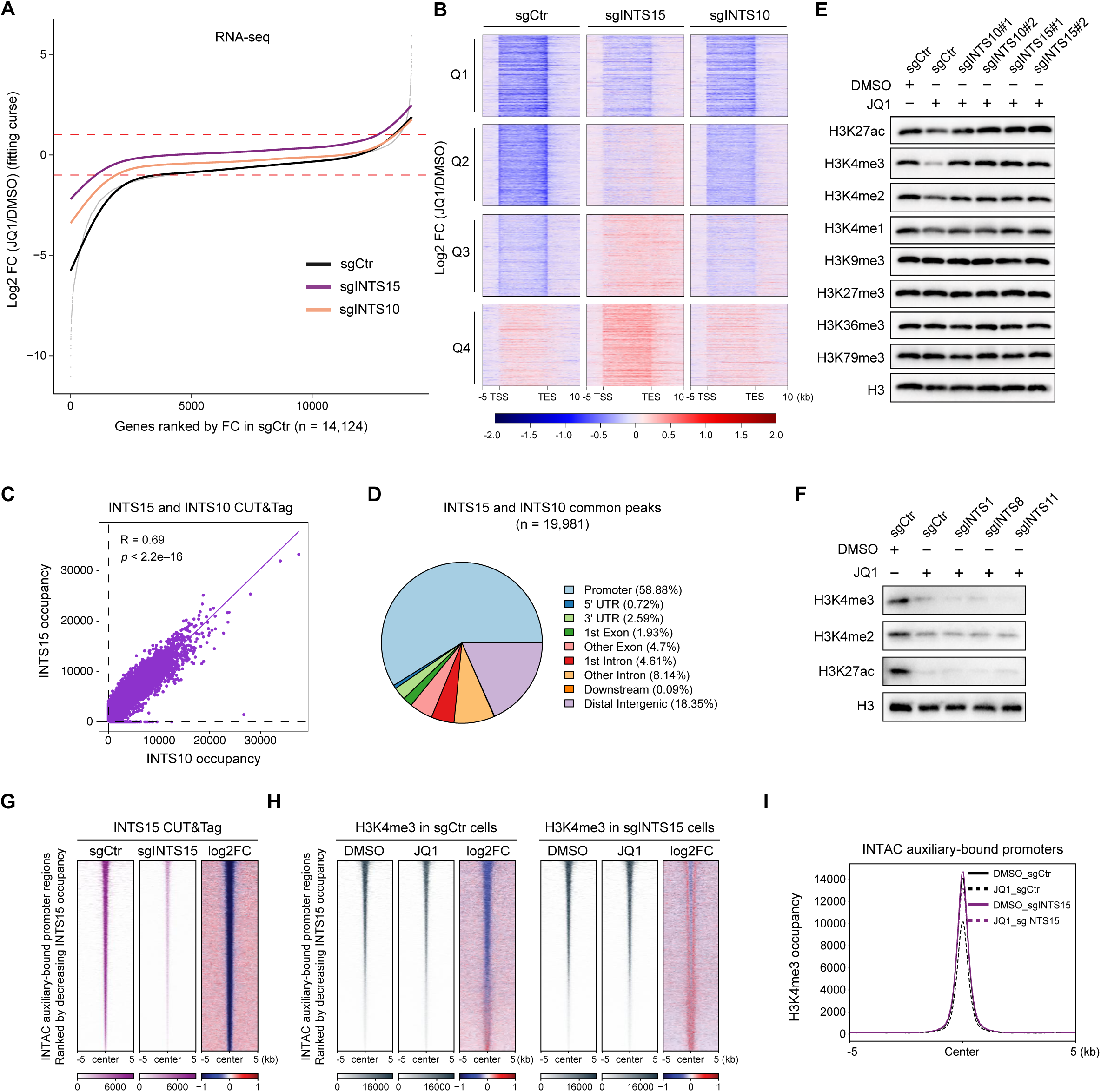
The INTAC auxiliary/arm module have broad roles in regulating gene expression and epigenetic landscape. (A) Fold change of gene expression (JQ1 versus DMSO_sgCtr) in sgCtr, sgINTS15 and sgINTS10 cells against JQ1 treatment, genes ranked by increasing FC in sgCtr cells. (B) Heatmaps for fold change of gene expression (JQ1 versus DMSO_sgCtr) in sgCtr, sgINTS15 and sgINTS10 cells, genes equally quartalized into four clusters (Q1, Q2, Q3 and Q4) and ranked by increasing FC in sgCtr. (C) Two-dimensional density plot showing the genome-wide occupancy of INTS10 (x axis) and INTS15 (y axis). (D) Pie chart showing the genomic distribution of the common peaks of INTS10 and INTS15. (E) Western blotting on histone modifications in sgCtr, sgINTS10 and sgINTS15 cells treated with DMSO or JQ1 for 48 hours. H3 serves as a loading control. (F) Western blotting on histone modifications in sgCtr, sgINTS1, sgINTS8 and sgINTS11 cells treated with DMSO or JQ1 for 48 hours. H3 serves as a loading control. (G) Heatmaps of INTS15 occupancy [reads per million mapped reads (RPM) per bp and log_2_ fold change (FC)] measured by CUT&Tag in sgCtr and sgINTS15 cells, ranked by decreasing INTS15 occupancy in sgCtr cells. (H) Heatmaps showing the H3K4me3 occupancy (reads per million mapped reads (RPM) per bp and log2 fold change (FC)) in DMSO or JQ1 treated sgCtr (Left) and sgINTS15 cells (Right), ranked by decreasing INTS15 occupancy in sgCtr cells. (I) Metaplots showing the average levels of H3K4me3 centered at INTAC auxiliary module-bound promoters in sgCtr and sgINTS15 cells post DMSO or JQ1 treatment. See also Figure S3.

As a frequently amplified proto-oncogene in cancer, MYC is arguably the most well-known target of BET proteins and also a global activator of transcription.^14,18,84,85^ Nonetheless, in Eμ-Myc lymphoma cells, MYC is ectopically expressed and remained unaffected by JQ1 treatment (Fig. S3E–S3F). Moreover, the depletion of the INTAC auxiliary did not alter MYC expression levels, ruling out the possibility that the INTAC auxiliary’ regulation of BET inhibition sensitivity operates through MYC (Fig. S3E–S3F). To ascertain whether the INTAC auxiliary directly targets specific genes, we employed Cleavage Under Targets and Tagmentation (CUT&Tag)^86^ for INTS15 and INTS10 to delineate their genomic distributions. The analysis revealed a significant overlap between INTS15 and INTS10 binding sites, with a proportional occupancy across these regions (Fig. 3C and S3G). Identifying these regions co-occupied by INTS15 and INTS10 as targets of the INTAC auxiliary, we analyzed their genomic distribution, finding a predominant targeting of promoter regions (Fig. 3D). This implies the specific role of INTAC auxiliary in targeting gene promoters to regulate transcription under BET inhibition.

### The INTAC auxiliary/arm module regulates epigenetic landscape

Given that BET proteins recognize and regulate acetylated histones, we posited that these epigenetic marks might play a role in both the transcriptional repression induced by JQ1 and the derepression resulting from depletion of the INTAC auxiliary. To investigate this, we profiled the bulk levels of various transcription-related histone modifications following JQ1 addition. As anticipated, we observed a decrease in H3K27ac. Moreover, JQ1 treatment also led to a profound decline in the levels of H3K4me3 and a reduction in H3K4me2, with no noticeable changes in other modifications we examined (Fig. 3E).

Importantly, the diminished levels of histone marks, including H3K4me3, were reversed to normal upon the loss of INTS15 and INTS10 (Fig. 3E). Notably, depleting subunits of other INTAC modules did not rescue the decline of H3K4 methylation and histone acetylation, consistent with our prior observation that the auxiliary subunits regulate the BET inhibition sensitivity independently of the two catalytic activities of INTAC (Fig. 3F). To determine whether the regulation of H3K4 methylation by JQ1 and INTAC auxiliary was specific for Eμ-Myc cells, we treated parental or INTS15-depleted THP-1 and HGC-27 cells, both were sensitive to JQ1 and gained resistance upon the depletion of the INTAC auxiliary (Fig. S1G–S1H). Consistent with Eμ-Myc cell outcomes, JQ1 led to reduced H3K4 methylation levels, which were subsequently restored with the loss of INTS15 (Fig. S3H–S3I). However, the elimination of INTS15 did not mitigate the downregulation of MYC protein induced by JQ1 in these cell types (Fig. S3H–S3I). Given these finding, and considering the enhanced sensitivity to JQ1-induced proliferation inhibition with the presence of the INTAC auxiliary (Fig. S1G–S1H), we conclude that the INTAC auxiliary module confers BET inhibition sensitivity through mechanisms independent of MYC protein expression.

To further confirm JQ1-induced alterations of H3K4 methylation, we conducted H3K4me3 CUT&Tag in both control and INTS15-depleted cells. Most of genes downregulated by JQ1 exhibited declined H3K4me3 occupancy at their promoters, in line with the promoting role of H3K4me3 in transcription activation (Fig. S3J). In all genomic regions or promoters bound by the INTAC auxiliary (Fig. 3F and S3K), JQ1 treatment led to a widespread reduction in H3K4me3 levels, particularly at loci with high occupancy of both INTAC auxiliary and H3K4me3 (Fig. 3G and S3L). Notably, INTS15 depletion effectively negated the JQ1-induced decrease in H3K4me3 at these INTAC auxiliary target sites (Fig. 3H–3I and S3L–S3M). These findings unveiling the pervasive role of the INTAC auxiliary in regulating genome-wide epigenetic landscape.

### The INTAC auxiliary/arm module specifically interacts with complexes of epigenetic regulators

To elucidate the mechanisms of INTAC auxiliary-mediated regulation of histone marks, we performed protein purifications in cells overexpressing individual components the INTAC complex: INTS15 and INTS10 for the auxiliary module, and INTS6 and INTS11 to represent the phosphatase and RNA endonuclease modules, respectively. Mass spectrometry analysis of these purifications retrieved components of different INTAC modules in a stoichiometric manner, supporting the notion that INTAC assembly is highly modular within cells (Fig. S4A–S4D).^67^ To identify specific interactors of the auxiliary module distinct from other INTAC modules, we analyzed the proteins co-purified with INTS15/10 in comparison with those with INTS6/11. Notably, INTS15 and INTS10 exhibited not only a co-enrichment with other auxiliary subunits but also stronger interactions with multiple subunits of several chromatin-bound complexes, including RACK7/ZMYND8–KDM5C, NuRD, INO80, and LEC (Fig. 4A–4D). Considering that the H3K4 demethylase activity of RACK7–KDM5C and histone deacetylase function of NuRD could potentially account for the observed decreases in H3K4 methylation and H3K27ac, we confirmed their interactions with INTAC by co-immunoprecipitation (co-IP) analysis, with western blotting validating a preferred association of RACK7–KDM5C and NuRD with INTAC auxiliary subunits (Fig. 4E).

**Figure 4.**
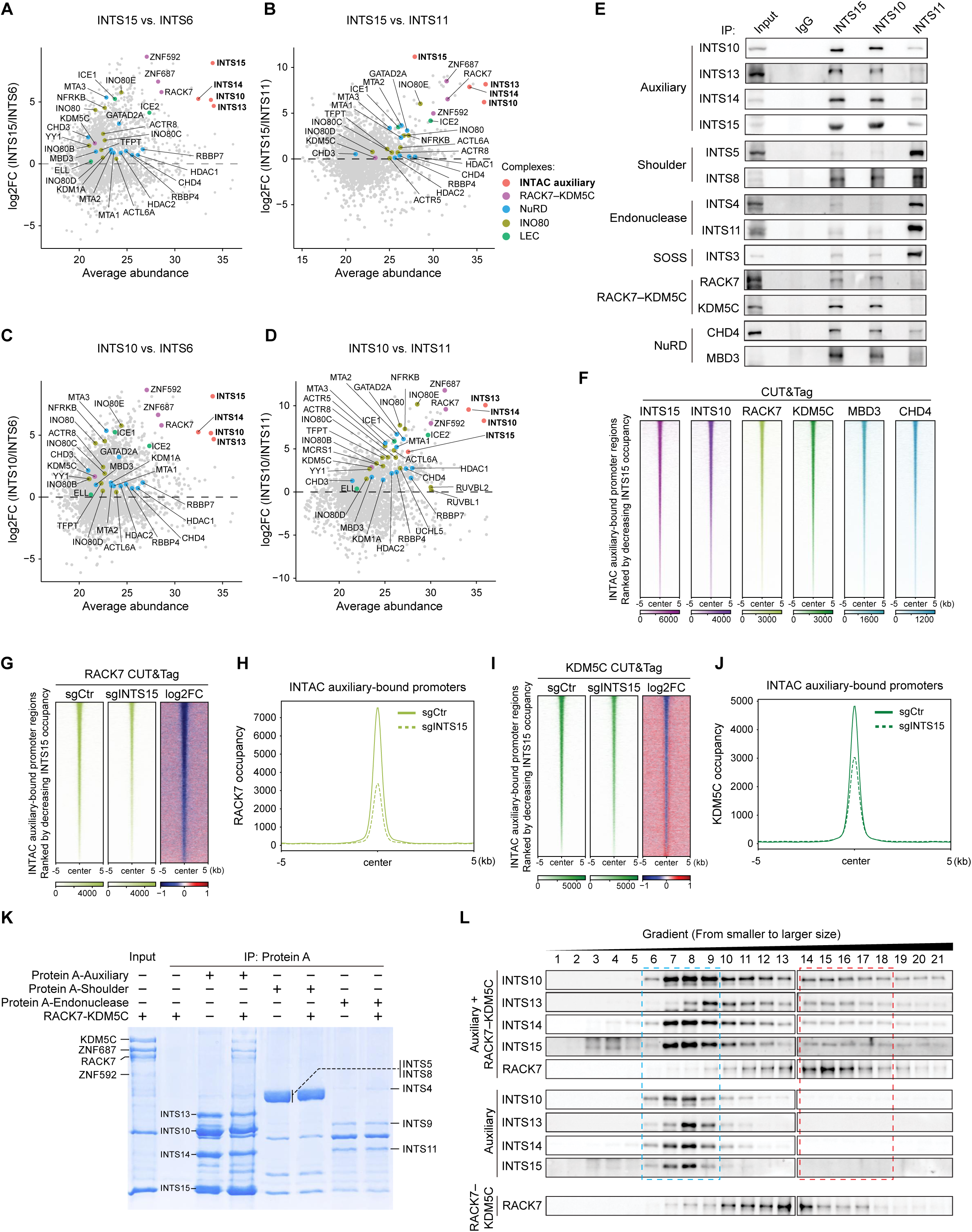
The INTAC auxiliary/arm module directly interacts with and facilitates the recruitment of the RACK7–KDM5C complex. (A-D) Volcano plots displaying proteins identified by mass spectrometry of Flag-labeled INTS15, INTS10, INTS6 and INTS11 over Flag-vector control. Quantification of log2 fold change (FC) for abundant binding proteins of INTS15 versus INTS6 (A), INTS15 versus INTS11 (B), INTS10 versus INTS6 (C) and INTS10 versus INTS11 (D). (E) Co-IP of endogenous INTS15, INTS10 and INTS11, then western blotting with the antibodies indicated. IgG serves as a negative control. (F) Heatmaps showing INTS15, INTS10, RACK7, KDM5C, MBD3 and CHD4 occupancy at INTAC auxiliary module-bound promoters, ranked by decreasing occupancy of INTS15. (G) Heatmaps showing RACK7 occupancy in sgCtr, sgINTS15 cells and log_2_FC (sgINTS15 versus sgCtr) at INTAC auxiliary module-bound promoters, ranked by decreasing occupancy of INTS15. (H) Metaplots showing the average levels of RACK7 occupancy in sgCtr and sgINTS15 cells centered at INTAC auxiliary module-bound promoters with H3K4me3. (I) Heatmaps showing KDM5C occupancy in sgCtr, sgINTS15 cells and log_2_FC (sgINTS15 versus sgCtr) at INTAC auxiliary module-bound promoters, ranked by decreasing occupancy of INTS15. (J) Metaplots showing the average levels of KDM5C occupancy in sgCtr and sgINTS15 cells centered at INTAC auxiliary module-bound promoters with H3K4me3. (K) In vitro pull-down assays using immobilized the auxiliary, shoulder, and RNA endonuclease modules of INTAC as baits incubated with RACK7-KDM5C. The bound proteins were subjected to SDS-PAGE followed by Coomassie blue staining. (L) Gradient centrifugation of the purified auxiliary module of INTAC incubated with RACK7-KDM5C (top), INTAC auxiliary module alone (median) and RACK7-KDM5C alone (bottom). Fractions were subjected to SDS-PAGE followed by western blotting. See also Figure S4.

Therefore, we proposed that the INTAC auxiliary might modulate H3K4 methylation and H3K27ac levels by facilitating the recruitment of RACK7–KDM5C and NuRD to chromatin. To examine it, we conducted CUT&Tag for RACK7, KDM5C, and two NuRD subunits (MBD3 and CHD4) in both control and INTS15-depleted cells. The genomic distribution of RACK7–KDM5C and NuRD largely overlapped with that of the INTAC auxiliary subunits (Fig. S4E–S4F), with the combined analysis identifying 13,065 loci co-bound by all three complexes (Fig. S4G). Intriguingly, INTS15 depletion resulted in marked decline in the occupancy of RACK7 and KDM5C at INTAC auxiliary-bound promoters (Fig. 4G–4J), while NuRD occupancy increased at these regions (S4H–S4K). The differential impacts on RACK7– KDM5C and NuRD occupancy were consistently observed across all INTAC auxiliary-associated peaks (S4L–S4O). These data reveal a preferred association of the INTAC auxiliary with multiple chromatin-bound complexes, indicating that the optimal recruitment of RACK7–KDM5C is contingent upon the presence of the auxiliary module.

### Direct interaction between the INTAC auxiliary/arm module and RACK7/ZMYND8–KDM5C

The observed dependence of INTS15 for the recruitment of RACK7–KDM5C to chromatin prompted us to delineate the molecular details of their interaction. To confirm direct physical contact, we reconstituted the auxiliary, shoulder, and RNA endonuclease modules of INTAC along with the RACK7–KDM5C complex for in vitro pulldown assays. Consistent with prior purification and co-IP results, a direct interaction was observed exclusively between the INTAC auxiliary module and RACK7–KDM5C (Fig. 4K). To further examine the strength of their interaction, we performed gradient centrifugation with the INTAC auxiliary module either alone or after incubation with RACK7–KDM5C. Typically, auxiliary subunits were predominantly found in fractions #6 to #9 (Fig. 4L, blue box), but co-incubation led to a shift of some auxiliary components towards denser fractions (#14 to #18), which also contained RACK7– KDM5C (Fig. 4L, red box). Moreover, the presence of INTAC auxiliary caused RACK7 to migrate into higher-molecular-weight fractions (Fig. 4L). These results unequivocally demonstrate a direct interaction between the INTAC auxiliary module and the RACK7–KDM5C complex.

### RACK7/ZMYND8–KDM5C mediates JQ1-induced decline in H3K4 methylation

To evaluate the implication of RACK7–KDM5C in JQ1-induced downregulation of H3K4 methylation, we blocked catalytic activity of KDM5C with its inhibitor KDM5-C70 alongside JQ1 treatment. Notably, treatment with KDM5-C70 dose-dependently restored H3K4me3 and H3K4me2 levels, either completely or partially, to their normal states (Fig. 5A). In line with KDM5C inhibition, the deletion of RACK7 by two distinct sgRNAs effectively prevented JQ1-triggered decline in H3K4 methylation (Fig. 5B and S5A). In contrast, the depletion of MBD2 and MBD3—two NuRD subunits crucial for the complex assembly and chromatin recruitment—did not reverse JQ1-induced diminishment of H3K4 methylation levels (Fig. S5B–S5C).

**Figure 5.**
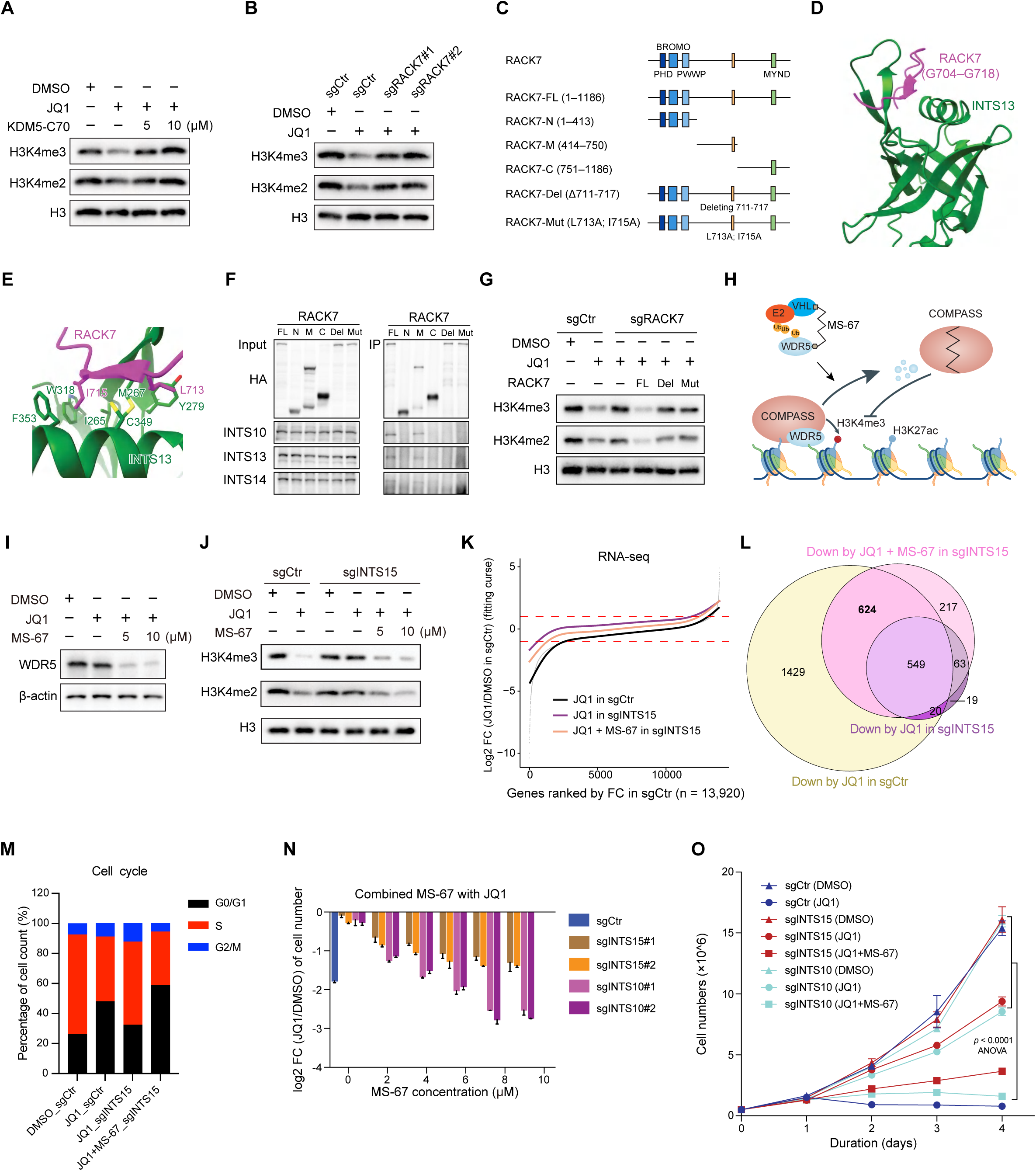
WDR5 degradation-induced diminishment of H3K4 methylation resensitizes the resistant cells to BET inhibition. (A) Western blotting for H3K4me3 and H3K4me2 levels in Eμ-Myc cells under conditions of DMSO, JQ1, combined WDR5 degrader MS-67 with JQ1. H3 is served as a loading control. (B) Western blotting for H3K4me3 and H3K4me2 levels in sgCtr and sgRACK7 cells post DMSO or JQ1 treatment. (C) Cartoon models showing the RACK7 truncations and mutants designed for exploring its interaction with INTAC auxiliary module. (D and E) The physical contact surface (C) and amino acids (D) between RACK7 and INTAC auxiliary module based on AlphaFold-Multimer (AF2-multimer) prediction. (F) Co-IP of endogenous INTS15, INTS10 and INTS11 with HA antibody in HA-tagged RACK7 truncations and mutants expressed cells. (G) Western blotting for H3K4me3 and H3K4me2 in sgCtr, sgRACK7 and ectopic RACK7-full length, - deletion and point mutants in sgRACK7 cells upon JQ1 treatment. (H) Cartoon models depicting the disruption of COMPASS via MS-67, a degrader targeting WDR5. (I) Western blotting for WDR5 post DMSO, JQ1 or MS67 treatment in Eμ-Myc cells. (J) Western blotting for H3K4me3 and H3K4me2 in sgCtr and sgINTS15 cells post DMSO, JQ1, or MS-67 combined with JQ1 treatment. (K) Curve plots showing transcriptional changes under the treatment of JQ1 in sgCtr and sgINTS15 cells, and MS-67 combined with JQ1 in sgINTS15 cells, respectively compared to expression in DMSO treated sgCtr cells, genes ranked by increasing FC in sgCtr cells. (L) Venn diagram showing overlaps for downregulated genes in sgCtr and sgINTS15 cells post JQ1 or MS-67 combined with JQ1 treatments. (M) Stacked plot showing the proportions of sgCtrl and sgINTS15 cells across different phases of cell cycle post DMSO, JQ1 or MS-67 combined with JQ1 treatment. (N) Cell survival assays of sgCtr, sgINTS15 and sgINTS10 cells under JQ1 or dosages of MS67 combined with JQ1 treatments. (O) Comparison on proliferation of sgCtr, sgINTS15 and sgINTS10 cells post JQ1 or MS67 combined with JQ1 treatments. See also Figure S5.

In pursuit of a RACK7 mutant defective in interacting with the INTAC auxiliary, we divided RACK7 into three segments and utilized AlphaFold-Multimer (AF2-multimer) to predict each segment’s interaction with INTAC auxiliary subunits (Fig. 5C). One such prediction indicated a β-strand within the middle region of RACK7 (RACK7–M) forming multiple interactions with INTS13 to form a β-sheet, with leucine 713 (L713) and isoleucine 715 (I715) being pivotal for the RACK7–INTS13 interaction (Fig. 5D– 5E). Immunoprecipitation results confirmed that only RACK7’s middle region interacted with the INTAC auxiliary module (Fig. 5F). Moreover, deleting the 7-residual β-strand, or mutating L713 and I715 to alanine, of RACK7 disrupted its association with INTAC auxiliary subunits (Fig. 5C and 5F). Subsequent rescue experiments by overexpressing either wild-type RACK7 or the mutants lacking INTAC auxiliary binding capability revealed that only the wild-type RACK7 overexpression sensitized RACK7-depleted cells to the JQ1-induced reduction in H3K4 methylation (Fig. 5G). These results indicate that the interaction of the INTAC auxiliary module and RACK7–KDM5C is crucial for determining the cellular responsiveness to JQ1-induced epigenetic alterations.

### The ablation of H3K4 methylation sensitizes the cells to BET inhibition

Considering the critical role of H3K4 methylation in determining the cellular response to BET inhibition, we posited that eradicating this methylation could shift INTAC auxiliary-depleted cells from resistance to sensitivity to JQ1 treatment. To test this hypothesis, we employed MS-67, a small molecule that specifically degrades WDR5—an integral component of the COMPASS complex essential for H3K4 methylation (Fig. 5H).^43,87^ Treatment with MS-67 resulted in a marked reduction in WDR5 levels (Fig. 5I). We thus treated INTS15-depleted cells with MS-67 and observed an apparent decline in H3K4 methylation under the condition of BET inhibition (Fig. 5J).

We further explored the impact of MS-67 on gene expression in conjunction with JQ1. The depletion of INTS15 had previously been shown to diminish the transcriptional suppression effect of JQ1; however, when we additionally introduced MS-67 to this condition, we observed a significant reinstatement of cellular sensitivity to JQ1’s effect on gene expression (Fig. 5K and S5D–S5F). Most genes downregulated by the combined MS-67 and JQ1 treatment in INTS15-deficient cells were overlapped with those affected by JQ1 in control cells (Fig. 5L), raising the possibility that the dual inhibition by MS-67 and JQ1 could induce cellular responses in INTS15-depleted cells that are similar to those observed with JQ1 treatment in control cells. Indeed, our cell cycle analysis indicated that although the absence of INTS15 mitigated JQ1’s induction of cell cycle arrest, the joint application of MS-67 and JQ1 prompted a significant G1 phase arrest in INTS15-depleted cells, similar to the arrest observed in control cells treated with JQ1 (Fig. 5M and S5G). Furthermore, MS-67 treatment increased the vulnerability of previously resistant INTS15-depleted cells to the growth-inhibitory effects of JQ1 in a dose- and time-dependent manner (Fig. 5N-O). These results support the strategy of ablating H3K4 methylation through the targeted degradation of WDR5 as a promising tactic to overcome resistance to BET inhibition.

## DISCUSSION

Our study has utilized genome-wide CRISPR screens to explore the mechanisms of the antiproliferative effect of BET inhibitors and the basis of intrinsic and acquired resistance to BET-targeted therapy. These screens not only support prior studies^78^ reporting the involvement of the autophagy pathway in BET-inhibition-induced cell death but have also revealed a previously unappreciated but significant contribution of the auxiliary/arm module of the INTAC complex to the efficacy of BET inhibition. Importantly, subunits of other INTAC modules were not enriched in our screens, nor did their individual disruption affect the sensitivity to BET inhibitors. Meanwhile, the loss of the INTAC auxiliary module had a minimal effect on snRNA processing, corroborating prior studies.^71–74^ Furthermore, our data showed that the INTAC auxiliary module did not influence the phosphatase activity of INTAC, aligning with structural analyses showing the auxiliary and phosphatase modules are located on opposite ends of the complex.^88,89^ These observations imply that while the auxiliary module can integrate with INTAC and support the functioning of other modules in specific developmental contexts,^71–74^ it likely holds independent roles beyond the two catalytic activities of the complex.

BET inhibition results in pronounced reduction in the levels of H3K27ac and H3K4me2/3, both of which are associated with active transcription, while gene body-related modifications, such as H3K36me3 and H3K79me3, along with repressive marks, including H3K9me3 and H3K27me3, remained unaffected as measured at the bulk level. Notably, JQ1-induced diminishment of these marks and associated gene expression necessitates the presence of the INTAC auxiliary module. Through analyses involving protein purification, endogenous co-immunoprecipitation, in vitro pulldown, and gradient centrifugation assays, we established a direct and stable interaction between the INTAC auxiliary module and the H3K4 demethylase complex RACK7–KDM5C. Depleting the INTAC auxiliary module resulted in a pervasive decrease in RACK7–KDM5C occupancy, suggesting the auxiliary module’s role in guiding the chromatin recruitment of RACK7–KDM5C.

Similar to the effects seen with the loss of auxiliary INTAC subunits, depletion of RACK7 or inhibition of KDM5 activity prevented the JQ1-triggered changes in H3K4 methylation, highlighting the pivotal role of the INTAC auxiliary–RACK7–KDM5C axis in mediating the removal of H3K4me2/3 in the context of BET inhibition (Fig. S5H–S5J). However, it is important to note that the RACK7–KDM5C complex comprises subunits equipped with various DNA-binding zinc finger motifs and domains that recognize specific histone marks, potentially facilitating its recruitment to chromatin.^56–59^ This complexity underscores the necessity for further research to unravel how these elements synergize with the INTAC auxiliary module, influencing the genomic localization of the RACK7–KDM5C complex and its functional dynamics in transcription regulation.

This study, along with two recent preprints,^88,89^ reveals that the INTAC auxiliary module forms specific interactions with multiple chromatin and transcription regulators, in addition to RACK7–KDM5C. Structural analyses from Sabath et al.^89^ and our AF2-multimer predictions identified a motif on RACK7 crucial for its interaction with the INTAC auxiliary module, which enabled us to engineer RACK7 mutants that disrupt this interaction. As anticipated, JQ1-induced reduction in H3K4 methylation was observed exclusively with the overexpression of wild-type RACK7 in RACK7 knockout cells, but not with the RACK7 point mutants, thereby underscoring the critical role of the INTAC auxiliary–RACK7– KDM5C axis in modulating BET inhibition sensitivity. Furthermore, the INTAC auxiliary interaction motif found on RACK7 is also shared by other interactors, such as INO80 and the Little Elongation Complex (LEC), suggesting a competitive binding scenario for access to the INTAC auxiliary module, potentially leading to the formation of mutually exclusive complexes.^89^ Therefore, future studies are warranted to dissect the distinct biological roles facilitated by the INTAC auxiliary module’s interactions with each of these complexes and transcription factors.

We found that BRD4, and not BRD2 or BRD3, is the critical mediator of JQ1-induced growth inhibition and the INTAC auxiliary-regulated BET inhibition sensitivity. Compared with BRD2 and BRD3, BRD4 uniquely contains a C-terminal motif that interacts with the positive transcription elongation factor b (P-TEFb). This interaction stimulates the kinase activity of P-TEFb towards its substrates such as the C-terminal domain of Pol II’s largest subunit RPB1 and the transcription elongation regulators DRB Sensitivity Inducing Factor (DSIF) and negative elongation factor (NELF), which are associated with the transitioning of paused Pol II into productive elongation.^17,90–94^ Consistent with this notion, our data revealed that JQ1 treatment significantly reduced gene expression, an effect that was substantially counteracted by the depletion of the INTAC auxiliary module.

Pause-release dynamics are governed not just by specific factors interacting directly with Pol II but are also significantly influenced by the positioning and modifications of nucleosomes. Notably, recent studies have illuminated the role of H3K4me2/3 in modulating the stability of Pol II pausing and its subsequent release.^51,52^ Therefore, the maintenance of H3K4 methylation by BET proteins adds another layer of complexity to its regulation of transcriptional pausing and release. Conversely, the diminishment of H3K4 methylation by RACK7–KDM5C, in coordination with the INTAC auxiliary, might induce transcriptional repression by altering Pol II pause-release dynamics. Supporting the role of H3K4 methylation in transcriptional activation, targeted degradation of the COMPASS subunit WDR5 effectively leads to significantly decreased H3K4 methylation and extensive transcriptional silencing. This action effectively resensitizes the previously BET resistant tumor cells with impairment of the INTAC auxiliary module (Fig. S5K). Collectively, our findings highlight the potential of simultaneously targeting coordinated chromatin and transcription regulators as a strategy to circumvent drug-resistant tumors by achieving substantial shifts in epigenomes and transcriptomes.

## Acknowledgments

This work was supported by grants from the National Key R&D Program of China (2021YFA1301700, 2021YFA1300100), the National Natural Science Foundation of China (32070636, 32300437).

## Author contributions

Conceptualization, F.X.C., H.J., and P.Z.; Experiments, P.F., X.-Y.S., and S.C. with help from Z.W., H.Z., B.T., W.X., W.J., H.Y., and C.X.; Sequencing and proteomics data analysis, A.S. with help from R.-Y.M. and J.C.; Writing, P.F., X.-Y.S., A.S., F.X.C with input from all authors; Supervision, F.X.C.

**Supplemental Figure Legends**

**Figure S1.**

(A) Table listing the top genes related to cell sensitivity to BET inhibition over 4-fold enriched through CRISPR screens. (B) Proliferation of sgCtr, sgINTS10, sgINTS13, sgINTS14 and sgINTS15 cells against JQ1 treatment. (C-F) The correlation between cell sensitivity to JQ1 and protein expression of INTAC auxiliary subunits INTS15 (C), INTS10 (D), INTS13 (E) and INTS14 (F). (G and H) Cell survival assays in sgCtr, sgINTS10, sgINTS13, sgINTS14 and sgINTS15 THP-1 (G) and HGC-27 cells (H) post dosages of JQ1 treatment.

**Figure S2.**

(A) Western blotting of the whole cell extraction in sgBRD2, sgBRD3 and sgBRD4 Eμ-Myc cells. (B) PCA analysis on transcriptomes of sgCtr, sgINTS10 and sgINTS15 cells under DMSO or JQ1 treatment. (C and D) MA plot illustrating the significantly up- and down-regulated genes in sgINTS15 (C) and sgINTS10 cells (D) post JQ1 treatment, in comparison with DEGs in sgCtrl cells against JQ1. (E) qPCR analysis for the representative genes that deficient auxiliary module restored against JQ1. (F) Flow cytometric evaluation of propidium iodide (PI) staining for cell-cycle analysis in sgCtr and sgINTS15 post DMSO or JQ1 treatment. (G) Cell survival assays of sgCtr, sgINTS15 and sgINTS10 cells under JQ1 or dosages of CDK4/6 inhibitor Abemaciclib combined with JQ1 treatments.

**Figure S3.**

(A-C) Curve plots showing gene expression (RPM per bp and log2 fold change) in JQ1 treated sgCtr cells (A), JQ1 treated sgINTS15 cells (B) and JQ1 treated sgINTS10 cells (C) versus DMSO treated sgCtr cells.

(A) Metagenes showing the log2 fold change of transcription levels on scaled genes in DMSO-and dTAG-treated sgCtr, sgINTS15 and sgINTS10 cells, quartalized into four clusters (Q1, Q2, Q3 and Q4) and ranked by increasing FC in sgCtr. (E) Western blotting for BRD2, BRD3, BRD4 and c-MYC in DMSO- and JQ1-treated sgCtr, sgINTS10 and sgINTS15 cells. β-actin is served as a loading control. (F) RNA-seq tracks at MYC gene in DMSO- and JQ1-treated sgCtr, sgINTS10 and sgINTS15 cells. (G) Venn diagram showing the overlap for CUT&Tag signals of INTS10 and INTS15. (H and I) Western blotting for H3K4me3, H3K4me2 and C-MYC in sgCtr, sgINTS10 and sgINTS15 THP-1 (H) and HGC-27 cells (I). (J) Pie chart showing H3K4me3 occupancy at promoters of down-regulated genes repressed by JQ1. (K) Heatmaps of INTS15 occupancy in sgCtr, sgINTS15 cells and log2FC (sgINTS15 versus sgCtr) at all INTAC auxiliary module-bound peaks, ranked by decreasing occupancy of INTS15. (L) Heatmaps showing the H3K4me3 occupancy (reads per million mapped reads (RPM) per bp and log2 fold change (FC)) in DMSO or JQ1 treated sgCtr (Left) and sgINTS15 cells (Right) at all INTAC auxiliary module-bound peaks, ranked by decreasing INTS15 occupancy in sgCtr cells. (M) Metaplots showing the average levels of H3K4me3 centered at all INTAC auxiliary module-bound peaks in sgCtr and sgINTS15 cells post DMSO or JQ1 treatment.

**Figure S4.**

(A-D) Volcano plots displaying proteins identified by mass spectrometry of Flag-labeled INTS15 (A), INTS10 (B), INTS6 (C) and INTS11 (D) over Flag-vector control. Components of different INTAC modules are labeled in red. (E and F) The overlaps for CUT&Tag signals of INTAC auxiliary module (INTS15 and INTS10) with RACK7-KDM5C (E) and CHD4 and MBD3 of NuRD complex (F). (G) Venn diagram showing the overlaps for genomic distribution of RACK7–KDM5C, NuRD and INTAC auxiliary components. (H and J) Heatmaps of MBD3 (H) and CHD4 occupancy (J) in sgCtr, sgINTS15 cells and log2FC (sgINTS15 versus sgCtr) at INTAC auxiliary module-bound promoters, ranked by decreasing occupancy of INTS15. (I and K) Metaplots showing the average levels of MBD3 (I) and CHD4 occupancy (K) in sgCtr and sgINTS15 cells centered at INTAC auxiliary module-bound promoters. (L-O) Metaplots showing the average levels of RACK7 (L), KDM5C (M), MBD3 (N) and CHD4 occupancy (O) in sgCtr and sgINTS15 cells centered at all INTAC auxiliary module-bound peaks.

**Figure S5.**

(A) Western blotting for RACK7 in sgCtr and sgRACK7 Eμ-Myc cells. (B and C) Western blotting for MBD2 and MBD3 (B), as well as histone marks (H3K4me3 and H3K4me2) in sgCtr, sgMBD2, sgMBD3 and double transfected (sgMBD2; sgMBD3) Eμ-Myc cells. β-actin and H3 are served as loading controls. (D-F) Curve plots showing gene expression (RPM per bp and log2 fold change) in JQ1 treated sgCtr cells (D), JQ1 treated sgINTS15 cells (E) and combined MS-67 with JQ1 treated sgINTS15 cells (F) versus DMSO treated sgCtr cells. (G) Flow cytometric evaluation of propidium iodide (PI) staining for cell-cycle analysis in DMSO treated sgCtr, JQ1 treated sgCtr, JQ1 treated sgINTS15 as well as combined MS-67 with JQ1 treated sgINTS15 cells. (H-K) Cartoon models illustrating the roles of INTAC auxiliary module as an epigenetic silencer conferring sensitivity to BET inhibition in cancer cells. The functions of INTAC auxiliary module are depicted in context of parental cancer cells (H), JQ1-sensitive cells (I), JQ1-resistant cells induced by the deficient auxiliary module (J), as well as re-sensitized cancer cells treated with MS-67 in basis of JQ1-resistant condition.

## METHODS

### Cell culture

Mouse Eμ-Myc p19^Arf-/-^ cells were cultured in B cell medium (45% Dulbecco’s modified Eagle’s medium and 45% Iscove’s modified Dulbecco’s media, supplemented with 10% fetal bovine serum, L-glutamate, and 50 μM β-mercaptoethanol) with 1× Penicillin-Streptomycin (Gibico). Human THP-1 and HGC-27 cells were cultured in RPMI 1640 medium supplemented with 10% fetal bovine serum (FBS, Yeasen), 1× Penicillin-Streptomycin. Human 293T cells were grown in DMEM supplemented with 10% FBS, 1× Penicillin-Streptomycin. All cells were cultured at 37°C, 5% CO_2_ and were negative for mycoplasma contamination.

### Genome-wide CRISPR screens

Mouse Two Plasmid Activity-Optimized CRISPR Knockout Library (Addgene#1000000096) was amplified in *E coli.* DH5α Electro-Cells according to the manufacturer recommendation. Eμ-Myc p19^Arf-/-^ cells were infected at a multiplicity of infection of approximately 0.3. One day after infection, medium was replaced with fresh growth medium. Two days after infection, cells were replated into growth medium containing puromycin. Cells were selected with puromycin for 2 days and then plated at a density of 0.5 million cells per milliliter, with treatment condition (DMSO/JQ1). After 3 days of selection with JQ1, cells were replated and allowed to recover for 3 days without JQ1 selection. Then cells were replated at a density of 0.5 million cells per milliliter, with treatment of JQ1 for another 3 days. At this point, the DMSO/JQ1-treated samples were collected and counted. The genomic DNA were extracted from the DMSO/JQ1-treated samples, and then amplified to be sequenced on Illumina NovaSeq instrument.

### RNA-seq

RNA was extracted from cells using TRIzol agent according to manufacturer protocol. Then the RNA was depleted rRNA with rRNA probes, and libraries were synthesized flowing steps, including fragmentation, first strand cDNA synthesis, second strand cDNA synthesis, end repair, adaptor ligation and PCR amplification. RNA-seq libraries were sequenced on Illunina Nova-seq instrument.

### Cell cycle analyses

After treated with drugs for 48 hours, cells were collected and fixed for one hour in 70% ethanol. Cells were then treated with 0.2% Triton X-100, 50 μg/ml propidium iodide and 100 μg/ml RNase A for one hour and analyzed by FACS.

### Immunoprecipitation and mass-spectrometry analysis

293T cells with overexpression of the flag-tagged full-length INTS10, INTS15, INTS6 and INTS11 protein were respectively resuspended and lysed in lysis buffer containing 20 mM Hepes (pH 7.4), 140 mM NaCl, 2 mM MgCl_2_, 10% glycerol, 0.5% NP-40, 0.2 mM EDTA, 2 mM DTT, 1× protease inhibitor for 30 min. The lysate was centrifuged at 16,000 rpm for 30 min, and the supernatant was collected and incubated with anti-flag M2 beads (sigma) for 2 hours at 4°C. Then, the beads were washed three times with lysis buffer and twice with wash buffer containing 20 mM Hepes (pH 7.4), 140 mM NaCl, 2 mM MgCl_2_, 1% glycerol, 0.02% NP-40, 0.2 mM EDTA. After washed five times as above, SDS loading buffer was directly mixed with the beads for Western blotting detection and the gels with proteins were cut to perform mass spectrometry.

### Protein expression and protein purification

The full-length INTS10, INTS13, INTS14 and INTS15 open reading frames were separately cloned into a modified pCAG vector, and INTS15 was tagged with an N-terminal Flag-4×Protein A. All plasmids were cotransfected to HEK Expi293 cells by polyethylenimine (Polysciences) at 37°C for 72 hours. Then, cells were harvested for lysis and purification. INTS5-INTS8 module, INTS4-INTS9-INTS11 module and RACK7-KDM5C complex were overexpressed and purified individually in a similar way. Overexpressed HEK Expi293 cells were resuspended and lysed in lysis buffer [50 mM Hepes (pH 7.4), 200 mM NaCl, 0.2% CHAPS, 5 mM MgCl_2_, 5 mM adenosine triphosphate (ATP), 10% glycerol, 2 mM dithiothreitol (DTT), 1 mM phenylmethylsulfonyl fluoride (PMSF), aprotinin (1 mg/ml), pepstatin (1 mg/ml), and leupeptin (1 mg/ml)] for 30 min at 4°C. The supernatant after centrifugation at 16,000 rpm for 30 min was incubated with immunoglobulin G (IgG) resins for 2 hours at 4°C. The resins were washed three times with lysis buffer and twice using wash buffer [50 mM Hepes (pH 7.4), 200 mM NaCl, 0.1% CHAPS, 10% glycerol, and 2 mM DTT]. The protein was cleaved for 2 hours and eluted out. The purified complex was used in in vitro assays and stored at −80°C.

### Pull-down assay

HEK Expi293 cells with INTS10-13-14-15 module, INTS5-8 module and INTS4-9-11 module overexpression were respectively resuspended and lysed in lysis buffer containing 30 mM Hepes (pH 7.4), 300 mM NaCl, 5 mM MgCl_2_, 5 mM ATP, 10% glycerol, 0.2% CHAPS, 2 mM DTT, 1 mM PMSF, aprotinin (1 mg/ml), pepstatin (1 mg/ml), and leupeptin (1 mg/ml) for 30 min. The lysate was centrifuged at 16,000 rpm for 30 min, and the supernatant was collected and adjusted to 150 mM NaCl concentration. The above extract was incubated with immunoglobulin G (IgG) resins for 2 hours at 4°C. Then, the resins were washed three times with lysis buffer and twice with wash buffer containing 30 mM Hepes (pH 7.4), 150 mM NaCl, 5 mM MgCl_2_, 5 mM ATP, 10% glycerol, 0.1% CHAPS, and 2 mM DTT. After incubating with RACK7-KDM5C complex for 2 hours at 4°C, the resins were washed five times as above. SDS loading buffer was directly mixed with the resins for western blotting.

### Density gradient centrifugation

In gradient centrifugation, the proteins of INTS10-13-14-15 module and RACK7-KDM5C complex were incubated for 2 hours at 4°C, and slowly layered on top of a 4-ml 10 to 40% (v/v) glycerol gradient (SW60 Ti) in buffer containing 20 mM Hepes (pH 7.4), 200 mM NaCl, 0.05% CHAPS, and 2 mM DTT and centrifuged at 34,000 rpm for 16 hours. Per 200 μl was sorted manually as a fraction from the top of the gradient tube and analyzed by western blotting.

### CUT&Tag

Cells were collected by centrifugation with twice washed by PBS. 5×10^5^ cells were typically used to obtain sufficient DNA extraction for library construction. The cells were centrifuged (600g, 3 min) at room temperature, washed twice with 800 μl of wash buffer (20 mM HEPES pH 7.5, 150 mM NaCl, 0.5 mM spermidine, 1× protease inhibitor) and finally resuspended with 100 μl of wash buffer in low-retention PCR tubes. The concanavalin-A-coated magnetic beads (Smart-Lifesciences) were activated in advance and resuspended with the same volume of the binding buffer (20 mM HEPES pH 7.5, 10 mM KCl, 1 mM CaCl_2_, 1 mM MnCl_2_). A total of 10 μl of activated concanavalin A beads was added to 5×10^5^ cells with incubation for 10 min under gentle rotation. The bead-bound cells were magnetized to remove the liquid with a pipettor and resuspended in 50 μl of antibody buffer (20 mM HEPES pH 7.5, 150 mM NaCl, 0.5 mM spermidine, 1× protease inhibitor, 0.05% digitonin, 0.01% NP-40, 2 mM EDTA). Next, 1 μl of antibodies were added to combine transcription factors or histone modification by rotating at 4 °C overnight. For the IgG control, Rabbit IgG was used instead. After successive incubation with mouse anti-rabbit IgG (Solarbio, 1:100 dilution) and rabbit anti-mouse IgG (Solarbio, 1:100 dilution) in 100 μl of antibody buffer for 1 h at room temperature, the bead-bound cells were washed three times with dig-wash buffer (antibody buffer without 2 mM EDTA) to remove the unbound antibody. The pA-Tn5 adapter complex was mixed in dig-300 buffer (20 mM HEPES pH 7.5, 300 mM NaCl, 0.5 mM spermidine, 1× protease inhibitors, 0.01% digitonin, 0.01% NP-40) to a final concentration of 0.2 μM. The bead-bound cells were resuspended in 100 μl of pA-Tn5 mix and incubated at room temperature for 1 h followed by removing the supernatant. After adequate washing, the tagmentation reaction was performed in 40 μl of tagmentation buffer (10 mM TAPS-KOH pH 8.3, 10 mM MgCl_2_, 1% DMF) at 37 °C for 1 h. Next, 1.5 μl of 0.5 M EDTA, 0.5 μl of 10% SDS and 1 μl of 20 mg/ml protease K were added to stop the reaction. After incubation for 1 h at 55 °C, DNA purification was performed using VAHTS DNA Clean Beads (Vazyme), and eluted in 10 μl of 0.1% Tween-20. The eluent was mixed with 10 U of Bst 2.0 WarmStart DNA polymerase (NEB) and 1 × Q5 polymerase reaction buffer (NEB) in a 20 μl reaction system. The reaction was completed at 65 °C for 30 min and then at 80 °C for 20 min to inactivate the Bst 2.0 WarmStart DNA polymerase. The purified DNA was amplified by Q5 high-fidelity DNA polymerase (NEB) with a universal i5 primer and a uniquely barcoded i7 primer. The exact PCR cycles were estimated by qPCR before amplification. PCR amplification with 13–14 cycles yielded enough quantity of library for sequencing. After library size-selection with 0.56–0.85 VAHTS DNA Clean Beads, with library sizes ranging from 200 to 700 bp, the products were next either analysed using qPCR or sequenced on the NovaSeq 6000 platform (Mingma Technologies).

## QUANTIFICATION AND STATISTICAL ANALYSIS

### RNA-seq data analysis

The paired raw reads were quality-checked with FastQC v0.11.9 (Babraham Institute) and trimmed by Trim Galore v0.6.6 (Babraham Institute) with the parameter “-q 25” to remove adaptors and low-quality reads. Then, the remaining reads were aligned to the mouse mm10 and human hg19 assemblies using STAR v2.7.5c^95^ with parameters “--outSAMtype BAM SortedByCoordinate –twopassModeBasic -- outFilterMismatchNmax 3”. Duplicates and low-quality reads were removed, and reads mapped in proper pairs were isolated using SAMtools v1.9^96^ with parameter “-f 2”. The number of spike-in hg19 reads was counted with SAMtools v1.9 and used to generate normalization factor alpha = 1e6/ hg19_count for coverage profiles. Strand-specific normalized bigwigs were generated. Raw gene counts were generated by featureCounts tool from the Rsubread R package v2.0.1.^97^ Differential expression analysis was performed using DESeq2 R package v1.26.0,^98^ and the counts for the human hg19 spike-in were used to estimate the size factors. Genes with false discovery rate < 0.05 and absolute log2FC > 1 were considered as differentially expressed genes.

### CUT&Tag data analysis

Raw CUT&Tag reads were trimmed using Trim Galore v.0.6.6 (Babraham Institute) in paired-end mode. Trimmed reads were aligned to mouse mm10 genome using Bowtie v.2.4.4 with the parameters “-N 1 -L 25 -X 700 --no-mixed --no-discordant”.^99^ Duplicated reads were removed with Picard Tools v.2.25.5 (Broad Institute) and the reads were shifted to compensate for the offset in tagmentation site relative to the Tn5 binding site using the alignmentSieve function of deepTools v.3.5.1 with the ‘--ATACshift’ option.^100^

### Statistical analyses

We used a Wilcoxon test throughout this study.

